# Deep learning-based prediction of tissue-specific splice sites in the human neural retina

**DOI:** 10.1101/2025.02.10.637427

**Authors:** Tabea V. Riepe, Suzanne E. de Bruijn, Susanne Roosing, Frans P.M. Cremers, Peter A.C. ‘t Hoen

## Abstract

Splice prediction tools can be used to identify splice-altering variants in patients with inherited diseases. Since splicing is tissue-specific, a predicted splice defect may vary depending on the tissue of interest. Current splice prediction tools often neglect splice junctions from the human neural retina, which can lead to the misinterpretation of benign and pathogenic variants in patients with inherited retinal diseases (IRDs). To address this issue, we developed a retina-specific splice prediction tool based on the model architecture of SpliceAI. In addition to retraining the existing model using splice junctions derived from retina RNA-sequencing data, we also improved the model’s generalization through hyperparameter optimizations such as dropout. The resulting retina-specific model was validated on retina-enriched exons and variants known to cause retina-specific splicing defects. Although the retina model correctly identified five more retina-enriched splice sites than the GTEx model, it did not enhance predictions for variants with tissue-specific splicing defects nor did it identify novel pathogenic variants in a set of genetically unexplained IRD cases. Despite these limitations, the retina-specific SpliceAI model shows promising results and should be applied to a larger patient cohort to uncover novel splice-altering variants in IRD patients.

## Introduction

Vision loss caused by inherited retinal diseases (IRDs) affects approximately 1 in 1,380 individuals [1]. Currently, only 52-74% of IRD patients receive a genetic diagnosis [2]. Various factors contribute to the significant number of unsolved IRD cases, and one factor is the oversight of genetic variants that alter mRNA splicing. Variants in canonical splice sites are easily detectable, but variants interfering with splicing that do not affect the AG/GT nucleotides flanking most exons are more difficult to interpret [3]. The effect of genetic variants on mRNA splicing can be assessed using reverse transcription polymerase chain reaction (RT-PCR), splice assays, or RNA-sequencing [4]. However, it is more effective to use computational tools called splice prediction tools to select likely splice-altering variants before performing experimental validation [5].

Several splice prediction tools are available [6–16], and recent advances in deep learning led to improvements in the field of splice prediction [17–19]. One main challenge of the currently available prediction tools is prediction of tissue-specific splicing. Tissue-specific splice prediction tools have been created but none of the tools were trained on splice junctions specific to the human neural retina [20–22]. This is a problem because splicing can be tissue-specific, and especially neural tissues like the retina are enriched for tissue-specific splicing [23,24]. Therefore, available splice prediction tools fail to predict the effect of some candidate IRD-causing variants on splicing correctly, demonstrating the need for a retina-specific splice prediction tool [25].

Previous studies have identified SpliceAI as one of the best splice prediction tools [26–28]. We trained modified versions of SpliceAI on human neural retina splice junctions extracted from publicly available RNA-sequencing data to create a retina-specific SpliceAI model that can provide an improved prediction of the effect of candidate IRD-variants.

## Methods

### Training data

The training data consists of RNA-sequencing data from 503 publicly available human retina samples from both healthy individuals and individuals suffering from age-related macular degeneration [29,30]. RNAs-sequencing data were trimmed with TrimGalore (v0.6.10) [31] and aligned to the ENSEMBL release 110 with STAR (v2.7.11a) [32] using two-pass alignment. The splice junction files from the different individuals were merged, and we prioritized canonical splice junctions with at least three uniquely mapped reads and a maximum spliced alignment overhang of at least five nucleotides (nt). We wrote a custom script to convert the STAR splice junctions to the required format for SpliceAI. In short, we selected junctions present in at least 15 samples and mapped the junctions to genes. Junctions not mapped to a known gene were excluded. We downloaded data from BioMart about paralogs of each gene and about transcription start sites (TSSs) and transcription termination sites (TTSs). We then selected splice sites for each gene that were located in between the first TSS and the last TTS for the gene. To reformat the STAR intron junctions to exon junctions, we subtracted one nt from the intron start sites and added one nt to the intron end sites. The dataset was split into training and testing by keeping splice junctions on chromosomes 1, 3, 5, 7, and 9 apart for testing. Additionally, genes that have a paralog were excluded from the test dataset due to sequence similarities with genes in the training dataset. In addition to the retina dataset, the GTEx and GENCODE V24lift37 splice junctions provided by Jaganathan *et al*. were used for comparison [6].

### Model architecture and training

As model architecture, we used the SpliceAI 32-layer deep neural network and trained it for 10 epochs unless stated otherwise. In addition to the standard SpliceAI model, several modifications including dropout, hyperparameter optimization, and fine-tuning were implemented. Dropout was included before the last convolution in each residual block and before the last convolution that generates the output, as proposed by Kim *et al*. [33]. We tested dropout rates of 0.2, 0.3, 0.4, and 0.5. For hyperparameter optimization, the encoding was replaced by label smoothing with labels [0.05, 0.05, 0.95] instead of [0, 0, 1], and the Adam optimizer was replaced by AdamW with a weight decay of 0.00001, as described by Zeng *et al*. [22]. The third modification included fine-tuning models trained on the GTEx dataset using the retina training data with different numbers of unfrozen layers (**Supplementary Figure S1A**). Additionally, we trained a model with the SpliceAI GTEx weights as initialization but without freezing any layers. For the fine-tuning, the number of epochs was reduced to five, and the learning rate was halved after every epoch.

### Testing of the models

Each model was tested on three different test datasets, the GENCODE and GTEx SpliceAI datasets, and the retina dataset created for this study. We evaluated the model using a threshold calibrated such that the number of predicted splice sites matched the number of actual splice sites in the test dataset. The performance measures calculated were top-k accuracy and the area under the precision-recall curve (PR-AUC). Top-k accuracy is the accuracy at the threshold for which the number of predicted splice sites matches the number of true splice sites in the test dataset.

### Predictions for retina-specific and control exons

For evaluation purposes, 115 splice acceptor sites and 107 splice donor sites not included in the retina or GTEx training data were extracted from 201 retina-enriched exons identified by Murphy *et al*. [34] and Ciampi *et al*. [35]. Murphy *et al*. defined retina-enriched exons as those with an inclusion level difference of at least 0.1 and a false discovery rate (FDR) of less than 0.01 between WT and Aipl1 knockout retinas [34]. Ciampi *et al*. identified retina-enriched exons using exon inclusion levels from *VastDB* [36] and the *Get_Tissue_Specific_AS*.*pl* [37] script. Additionally, we created a control dataset with non-tissue-specific splice sites that have similar properties to the splice acceptor and donor sites of the retina-enriched exons. Exons located on the same chromosome and strand as the retina-enriched exon with a similar length and GC content, were randomly selected. The cutoff used for retina-enriched splice sites was the optimal thresholds for the retina dataset defined by SpliceAI. For the control splice sites, the optimal threshold for the GTEx dataset was used.

### Predictions for variants with a retina-specific splicing defect

Four variants with a retina-specific splicing defect were selected from literature [25,38,39]. Delta scores for both the GTEx and retina SpliceAI models were calculated.

### Predictions for variants found in IRD patients

Whole genome sequencing (WGS) data were collected for 489 IRD patients. This cohort included both genetically explained and unexplained cases, originating from three sources: a 100-case IRD cohort from Fadaie *et al*. [40], a 100-case cohort with *USH2A*-associated retinitis pigmentosa from Reurink *et al*. [41], and additional unpublished studies focusing on genetically unexplained IRD cases. Genomic DNA was extracted from peripheral blood lymphocytes as previously described [42]. Sequencing was conducted by BGI on a BGISeq500 platform using either a 2x100 base pair (bp) or 2x150 bp paired-end module, or with the DNBseq Sequencing Technology, ensuring a minimum median genome coverage of 30-fold. Reads were mapped to the Human Reference Genome (GRCh38/hg38). Single-nucleotide variant (SNV) calling was performed using the Burrows-Wheeler Aligner (v.0.7814) [43] and Genome Analysis Toolkit HaplotypeCaller (Broad Institute). Structural variants (SVs) were identified using the Manta structural variant caller [44] and copy number variants (CNVs) were detected using Canvas Copy Number Variant Caller [45] based on read-depth analysis. Variants were then filtered for variants located in IRD-associated genes and present in five or less individuals. Additionally, the variants were annotated with VEP to add a gnomAD allele frequency to each variant. Only variants with an allele frequency < 0.05 and within 1,000 base pair (bp) distance of a transcript were prioritized for detailed analysis. Delta scores were calculated for both the GTEx and retina SpliceAI model and the variants were filtered for a delta score of 0.2 of higher for the retina model and a delta score below 0.2 for the GTEx model for the same splice site. Variants that passed the mentioned criteria were manually assessed to check the predicted splicing defect. For instance, variants predicted to strengthen an existing splice site or to weaken a non-existing splice site were excluded. Moreover, it was evaluated whether the affected gene was consistent with the patient’s phenotype

## Results

### Training data

The retina dataset derived from RNA-sequencing data contained 406,581 splice sites, of which 286,002 were located on the training chromosomes (**Table 1**). Most splice sites were detected in both RNA-sequencing studies, with only 7,032 sites missing from the Pinelli *et al*. dataset and 867 from the Ratnapriya *et al*. dataset. The number of splice junctions in the retina dataset was lower than the number of splice junctions in the GTEx dataset. However, the number was comparable to the number of splice junctions in the GENCODE dataset. Both the GTEx and GENCODE dataset were derived from the original SpliceAI model [6]. **Figure 1A** shows the overlap of splice junctions in the retina and GTEx datasets. About one-fourth of the retina splice junctions were unique to the retina, showing that many retina splice junctions are absent from the GTEx dataset. **Figure 1B** shows that 99% of the exons included in the retina training data had a canonical AG/GT splice motif, which was expected [46].

**Table 1:** Number of splice junctions in the GENCODE, GTEx, and retina dataset.

| Dataset | Splice junctions on all chromosomes | Splice junctions on training chromosomes (2,4,6,8,10-22,X,Y) |
| --- | --- | --- |
| GENCODE | 369,660 | 261,592 |
| GTEx | 556,174 | 391,515 |
| Retina | 406,579 | 286,002 |

**Figure 1:**
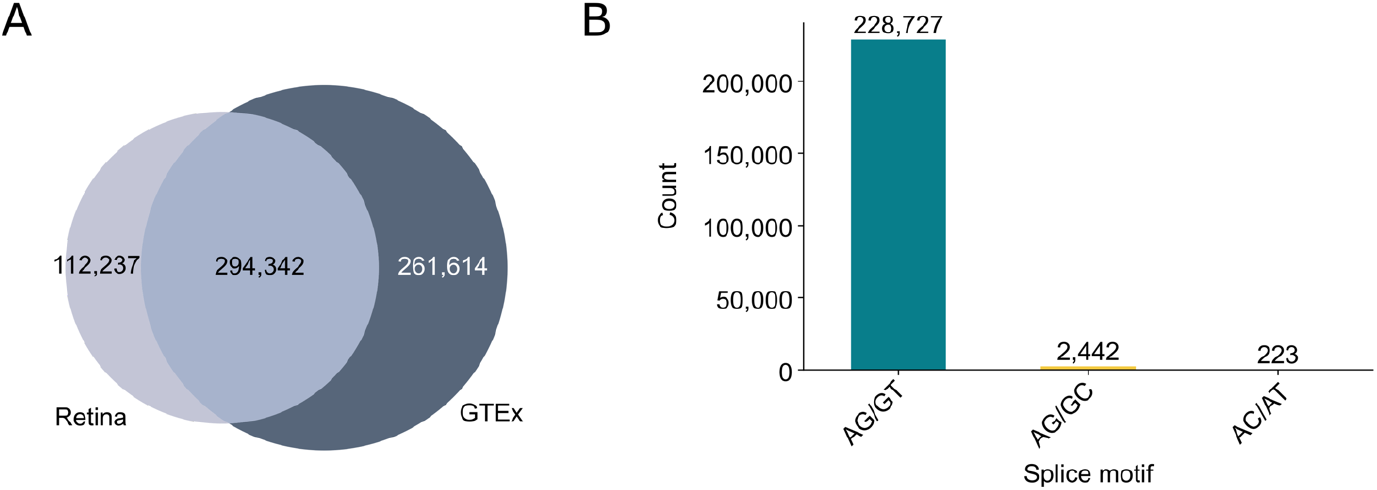
Characteristics of the retina training dataset. **(A)** Venn diagram comparing the splice junctions in the retina dataset (light grey) and GTEx dataset (dark grey). **(B)** Bar plot of the splice junction motif of the exons included in the retina training data.

### Comparison of retina and GTEx SpliceAI

We used the standard SpliceAI implementation to train two different models using identical parameters: one on the retina dataset and the other on the GTEx dataset. **Figure 2A** illustrates the PR-AUC for each model, displayed across training and validation datasets over each epoch. Both models achieved a PR-AUC of 0.9 or higher on their training datasets. However, the retina model, in particular, showed a notably lower PR-AUC on the validation dataset. This indicated that the retina model, and to a lesser extent the GTEx model, were experiencing overfitting. Both models were evaluated on the retina, GTEx, and GENCODE test dataset. For the retina test dataset, the retina model achieved PR-AUC of 0.81, while the PR-AUC for the GTEx model was 0.74 (**Table 2**). For the GTEx dataset, the GTEx model showed better performance with a PR-AUC of 0.87 compared to the PR-AUC of 0.83 of the retina model. Both models performed equally well on the GENCODE test dataset. Another difference between the two models was the optimal threshold. The retina model had a threshold of 0.22 for both acceptor and donor sites in the retina dataset and a threshold of 0.07 for both acceptor and donor sites in the GTEx dataset. For the GTEx dataset, the threshold was 0.39/0.38 for acceptor/donor sites in the retina dataset and 0.20/0.19 for acceptor/donor sites in the GTEx dataset. This showed that retina splice sites have a lower predicted score and were therefore probably weaker.

**Table 2:**
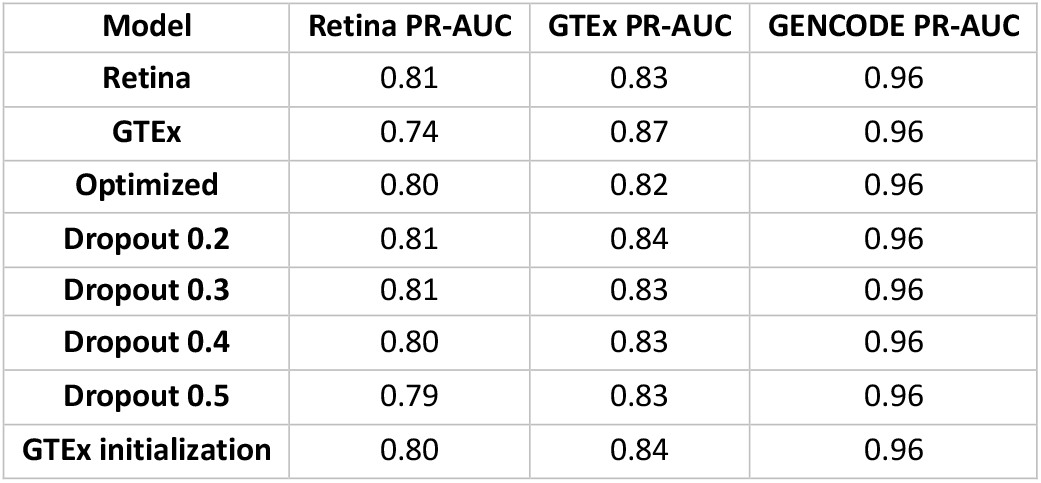
Comparison of the PR-AUC of the different models on the three test datasets.

**Figure 2:**
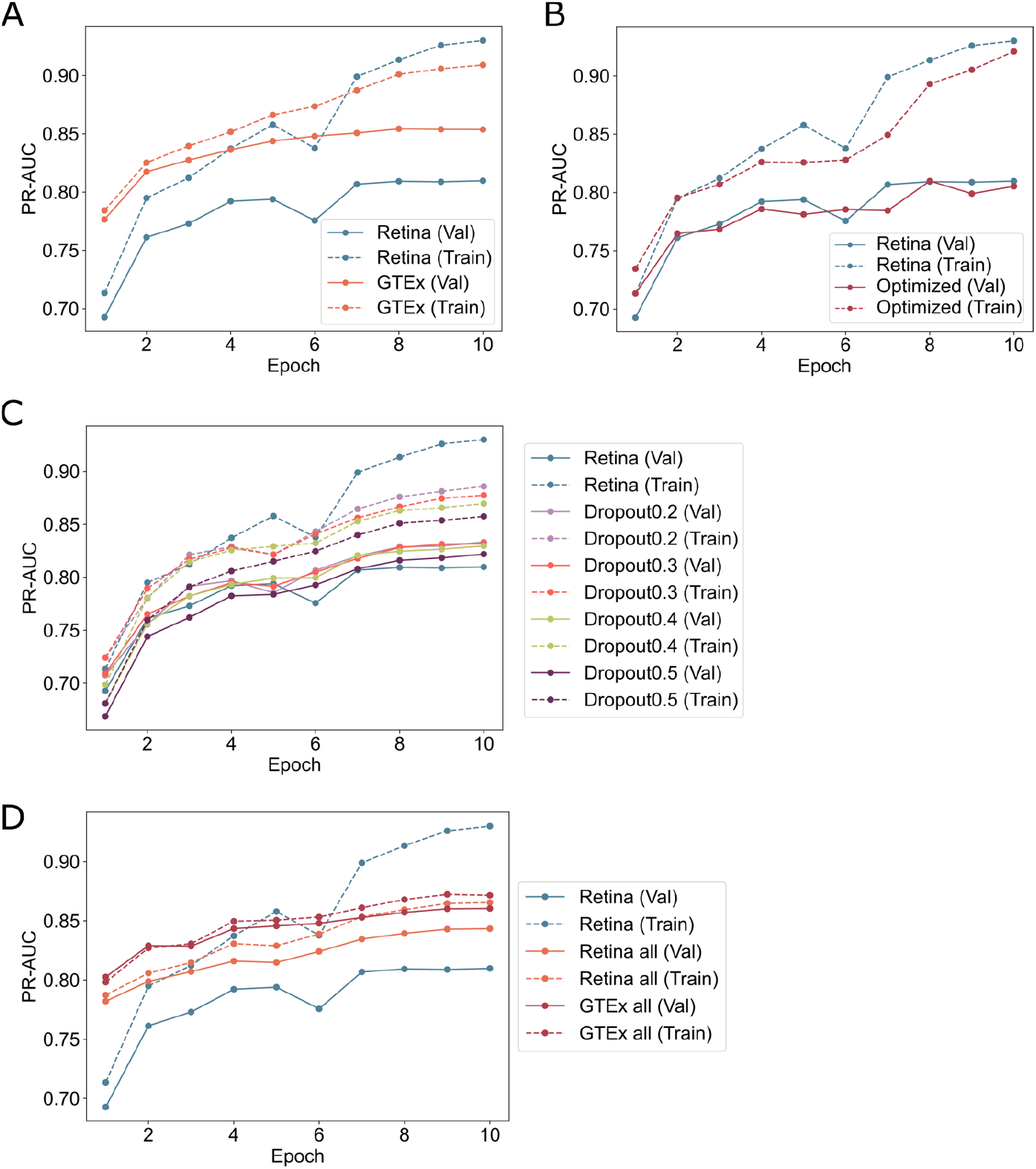
Average PR-AUC of the five models for each training epoch for different optimizations. **(A)** PR-AUC of the retina model (blue) and GTEx model (orange) on their respective training and validation datasets. **(B)** PR-AUC of the retina model (blue) and model with optimized parameters (red) for both the training and validation dataset. (**C)** PR-AUC of the retina model (blue) and dropout models for both the training and validation dataset. The four different dropout rates shown include 0.2 (lavender), 0.3 (orange), 0.4 (green), and 0.5 (purple). **(D)** PR-AUC of the retina model (orange) and GTEx model (red) trained on all chromosomes compared to the PR-AUC of the standard retina model (blue).

### Model with optimized parameters

Several variations of SpliceAI have been published [22,47–49]. Pangolin, for instance, uses different training settings like the AdamW optimizer instead of the Adam optimizer, label smoothing for better generalization and a warm-restart learning-rate schedule [22]. We incorporated the AdamW optimizer and label smoothing into the retina SpliceAI model. Including AdamW and label smoothing did not reduce overfitting, and the resulting model achieved a PR-AUC of 0.80 on the retina test dataset (**Figure 2B** and **Table 2**). The optimized parameters did not show an improvement compared to the standard retina model, and therefore, we decided to not include these settings in the final model.

### Dropout

SpliceAI is a deep convolutional network with many parameters, and it is prone to overfitting (**Figure 2A**). To prevent overfitting and increase the generalization of the model, we implemented a SpliceAI model with dropout and tested four different dropout rates of 0.2, 0.3, 0.4, and 0.5. All four dropout models showed a similar performance on the test datasets as the retina model (**Table 2**). We observed the lowest degree of overfitting for the model with a dropout rate 0.3 and therefore decided to use a dropout rate of 0.3 for our final model (**Figure 2C**).

### Transfer learning

Next, we applied a transfer learning approach by fine-tuning the previously trained GTEx model on the retina training data. The weights of the initial layers of the network were fixed, and solely the final layers were retrained. A summary of the six distinct configurations is provided in **Supplementary Figure S1**. Additionally, we trained a model starting with the GTEx initialization but without freezing weights. All the models with frozen weights had a PR-AUC comparable to the GTEx model, and only the model with no frozen weights showed an improved performance on the retina dataset (**Supplementary Figure S1**). Therefore, we chose to use the GTEx model weights as initialization for training the retina model but kept all layers unfrozen.

### Training using all chromosomes

To prevent overfitting and improve the retina SpliceAI model, we trained a model with a dropout rate of 0.3 using the GTEx model weights as initialization for ten epochs on splice junctions on all chromosomes. Additionally, we trained a GTEx model with the same settings. Both the GTEx and retina models trained on all chromosomes showed less overfitting and achieved a PR-AUC of about 0.85 on the training data (**Figure 2D**). These two models were used for the predictions in the rest of the study.

### Predictions for retina-enriched exons and control exons

To test if the retina model can predict retina-specific splice sites better than the GTEx model, both models were evaluated on a set of 115 retina-enriched acceptor sites and 107 retina-enriched donor sites. Additionally, we selected 115 control acceptor sites and 107 control donor sites. Most retina-enriched acceptor and donor sites with a prediction above the respective threshold were predicted by both the retina and GTEx models (**Figure 3A**). Nine sites were only predicted to be above the threshold by the GTEx model, and fourteen sites were only predicted by the retina model. The acceptor and donor sites in the control dataset were predicted to be above the threshold of both models, except for one donor site that was uniquely predicted by the GTEx model (**Figure 3B**).

**Figure 3:**
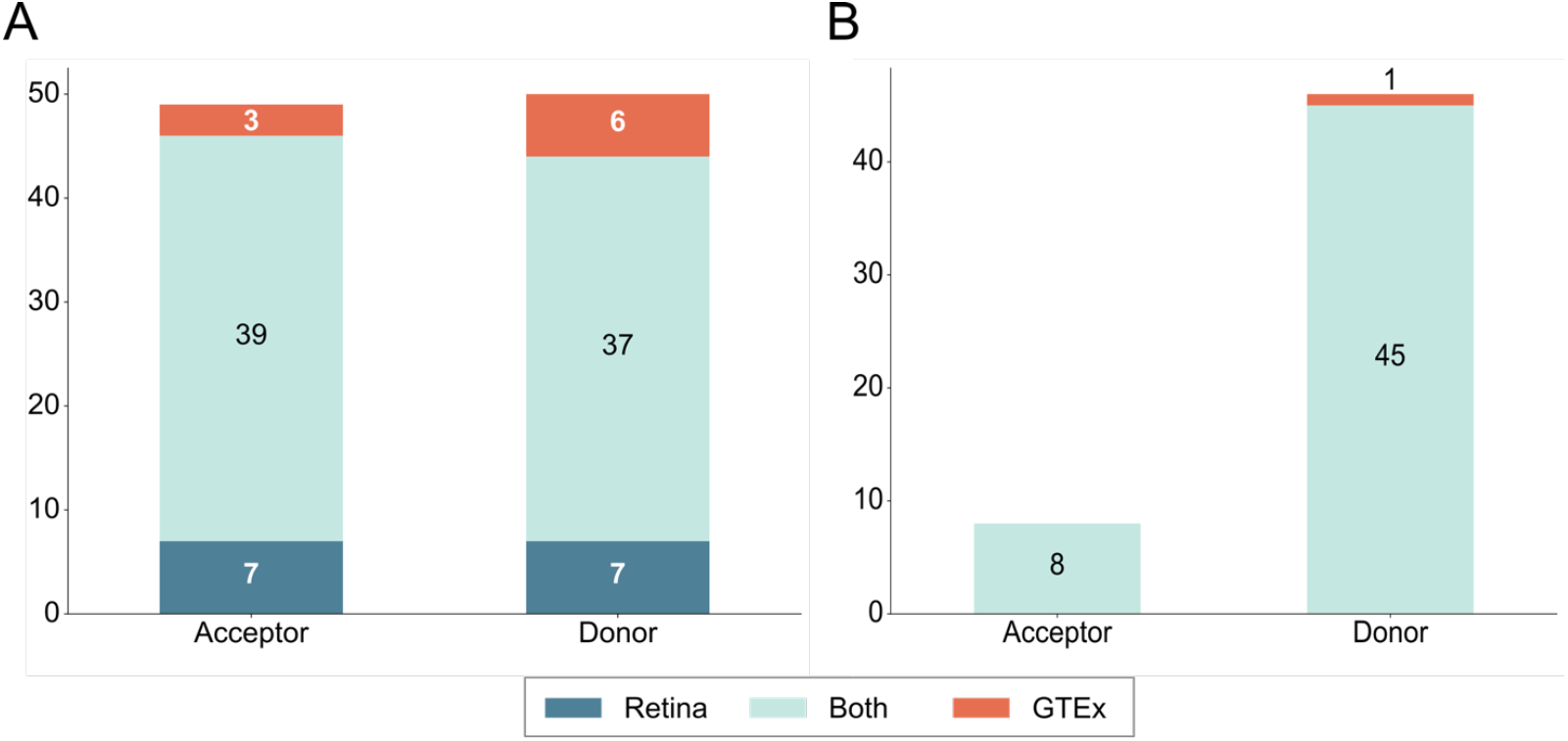
Retina and GTEx SpliceAI model predictions for retina-enriched and control exons. **(A)** Number of retina-enriched acceptor and donor splice sites that are predicted to be above the optimal threshold for the retina model (dark blue), the GTEx model (orange), or both models (light blue). **(B)** Number of control acceptor and donor sites that have a prediction above the threshold for either the retina model, the GTEx model, or both.

### Variants with a retina-specific splicing defect

We collected four variants with an experimentally validated retina-specific splicing defect from literature. Three of these variants were located in *ABCA4* and caused a pseudo-exon insertion between exon 30 and exon 31 by affecting retina-specific splicing enhancer and silencer motifs [25,50]. Additionally, the c.2991+1655A>G variant in *CEP290* is known to affect retina-specific splicing by activating a cryptic splice donor [38]. The predictions for the variants for both the retina and GTEx models can be seen in **Table 3**. The delta score of the c.4539+2001G>A variant for the acceptor gain was 0.10 for the retina model and 0.12 for the GTEx model. The delta score for the donor gain was 0.02 for the retina model and 0.05 for the GTEx model. All these scores were below the threshold of 0.2 that is commonly used to identify variants interfering with splicing. For the two other *ABCA4* variants, the scores for both acceptor and donor gain were 0.02 or lower. The *CEP290* variant had a predicted acceptor gain of 0.03 and a predicted donor gain of 0.14 for the GTEx model, which was higher than the scores of 0.01 and 0.02 predicted by the retina model. For all variants, both models predicted the acceptor and donor gain for the same position. While the acceptor gain position corresponded to the acceptor site of the *ABCA4* intron 30 pseudo-exon, the predicted donor site was different. Testing the retina and GTEx model on these variants with a retina-specific splicing defect showed that the GTEx model predicted higher delta scores for both the *ABCA4* c.4539+2001G>A and the *CEP290* variant.

**Table 3:** Retina and GTEx SpliceAI predictions for four variants with a retina-specific splicing defect. DS_AG = Delta score acceptor gain, DS_DG = Delta score donor gain, DP_AG = Delta position acceptor gain, DP_DG = Delta position donor gain

| Gene | Variant | Model | DS_AG | DS_DG | DP_AG | DP_DG |
| --- | --- | --- | --- | --- | --- | --- |
| ABCA4 | c.4539+2001G>A | Retina | 0.10 | 0.02 | 110 | -59 |
|  |  | GTEx | 0.12 | 0.05 | 110 | -59 |
|  | c.4539+2028C>T | Retina | 0.01 | 0.00 | 137 | -32 |
|  |  | GTEx | 0.02 | 0.01 | 137 | -32 |
|  | c.4539+1321A>G | Retina | 0.00 | 0.00 | -570 | 1321 |
|  |  | GTEx | 0.00 | 0.00 | -570 | 1321 |
| CEP290 | c.2991+1655A>G | Retina | 0.01 | 0.02 | 132 | 5 |
|  |  | GTEx | 0.03 | 0.14 | 132 | 5 |

### Predictions for variants found in IRD patients

The goal of this study was to create a retina-specific splice prediction tool that can be used to find novel splice altering variant in IRD patients. Therefore, we applied our retina SpliceAI model on a cohort of genetically explained and unexplained 489 IRD cases. Additionally, we generated predictions with the GTEx SpliceAI model trained by Jaganathan *et al*. [6]. We selected variants on 294 IRD-associated genes downloaded from RetNet [51] (19 May 2022). 86 of these variants had at least one predicted delta score of 0.2 or higher for retina SpliceAI and a predicted delta score lower than 0.2 for the same score with GTEx SpliceAI. Following assessment of the functional impact of the splicing defect and its correlation with the patient’s phenotype, four variants were identified as remaining candidates. (**Table 4**). For the patients with the *MKS1* and *RP1L1* variants, both associated with recessive inheritance, no second causative variant was identified in the WGS data for those genes. Additionally, the *RP1L1* variant was only supported by one read in the WGS data. The remaining two variants were identified in genes linked to dominant inheritance. The *PRPF31* variant matched the phenotype of the patient, but the variant did not co-segregate with the retinal phenotype in the family (absent in at least one affected family member). The *IMPDH1* variant was observed in a patient for who no dominant inheritance was reported by the clinician. We identified a second *IMPDH1* variant, c.1226G>A, classified as a variant of unknown significance by Franklin, in this patient and few literature reports suggest a potential link of *IMPDH1* variants and autosomal recessive inherited retinitis pigmentosa [52,53]. Additional analysis should be performed to check whether the two variants are located in trans or cis. Unfortunately, our analysis of the in-house IRD patient cohort did not result in a genetic diagnosis for unsolved IRD patients. However, further analysis and a larger cohort may increase the odds to find retina-specific splice defects.

**Table 4:** Summary of variants passing manual validation and corresponding to the patient’s phenotype. DS_AG = Delta score acceptor gain, DS_AL = Delta score acceptor loss, DS_DG = Delta score donor gain, DS_DL = Delta score donor loss, DP_AG = Delta position acceptor gain, DP_AL = Delta position acceptor loss, DP_DG = Delta position donor gain, DP_DL = Delta position donor loss

| Gene | Variant | DS_AG<br>Retina<br>(GTE<sub>x</sub>) | DS_AL<br>Retina<br>(GTE<sub>x</sub>) | DS_DG<br>Retina<br>(GTE<sub>x</sub>) | DS_DL<br>Retina<br>(GTE<sub>x</sub>) | DP_AG<br>Retina | DP_AL<br>Retina | DP_DG<br>Retina | DP_DL<br>Retina | Interpretation |
| --- | --- | --- | --- | --- | --- | --- | --- | --- | --- | --- |
| *IMPDH1* | NM_000883.4:<br>c.421_444del | 0.02<br>(0.02) | **0.43**<br>(0.05) | 0 (0.02) | **0.48**<br>(0.03) | 101 | -59 | -447 | 42 | Second variant:<br>NM_000883.4:c.1226G>A |
| *MKS1* | NM_017777.4:<br>c.515+12C>T | 0<br>(0.01) | **0.21**<br>(0.01) | 0.06<br>(0.11) | **0.2**<br>(0.04) | -2295 | 109 | 2 | 12 | No second variant |
| *PRPF31* | NM_015629.4:<br>c.946-408A>G | **0.24**<br>(0.1) | 0 (0) | **0.22**<br>(0.08) | 0<br>(0.02) | 0 | -19 | 75 | -1048 | Not found in affected<br>family member |
| *RP1L1* | NM_178857.6:<br>c.6361_6408dup | **0.27**<br>(0.02) | 0.07<br>(0) | **0.48**<br>(0) | 0.02<br>(0.01) | 0 | 110 | 0 | 66 | No second variant |

## Discussion

To account for tissue-specific splicing when predicting the effect of a variant on splicing, we developed a retina-specific version of SpliceAI. This model predicted more splice junctions in the retina dataset correctly while maintaining a good performance on the GTEx and GENCODE datasets. Moreover, the retina SpliceAI model identified more retina-enriched exons than the GTEx model but performed worse on a small set of variants with a retina-specific splicing defect.

Several modifications of SpliceAI have been published, and we implemented some of these for retina SpliceAI [22,47]. Zeng *et al*. proposed using label smoothing and the AdamW optimizer. However, applying these modifications to retina SpliceAI did not improve the performance of the model. One possible reason is that we did not optimize the hyperparameters for AdamW. Moreover, adding label smoothing likely introduced noise without improving the model’s generalization. Two modifications that did enhance the model were dropout and using the GTEx model weights as initialization.

A significant constraint of our study is the limited availability of retina-specific splicing data. Previous research identified retina-enriched exons and retina-specific splicing regulators like MSI1 and SRRM3 [34,35]. However, the exons are enriched in the retina but not exclusive to the retina, meaning the splicing motifs surrounding these exons are also recognized in other tissues. Additionally, retina-specific splicing events are rare and might not be adequately learned by the model. This could explain why the small set of three *ABCA4* intron 30 variants with experimentally confirmed retina-specific splicing demonstrated low delta scores for both the retina and GTEx models, as these variants interfere with retina-specific enhancer and silencer motifs. Moreover, only four variants are known to affect retina-specific splicing motifs, resulting in a very limited evaluation dataset. Most splice-altering variants are experimentally validated in splice assays in HEK293T cells, where retina-specific splicing effects would be overlooked.

SpliceAI, a convolutional neural network developed in 2019, has inspired attempts to create a splice prediction tool using a different method. For example, the nucleotide transformer showed similar performance for splice site predictions in the human genome as SpliceAI [54]. Other methods include ensemble predictors like SPiP or SQUIRLS [55,56], which combine predictions from several tools using a random forest classifier. Additionally, an explainable boosting classifier called AbSplice was developed [21]. AbSplice can consider tissue-specific splicing by using splice maps that exclude splice sites not used in the tissue of interest. A limitation of AbSplice for evaluating retina-specific splicing is that the retina is not included in GTEx, so only junctions not found in the retina can be excluded, and no novel retina-specific junctions can be added.

Next to validating already known retina-specific variants and acceptor and donor sites, we used retina SpliceAI to find novel splice-altering variants in IRD patients. Unfortunately, we did not identify any novel pathogenic variants. This could be due to the dataset consisting of WGS data from 489 patients, of which about 23% already received a conclusive genetic diagnosis [42]. Another reason might be that variants affecting retina-specific splicing contribute only a small number of unsolved IRD cases. Other potential causes for the lack of a genetic diagnosis include the presence of disease-causing variants in novel IRD genes, repetitive regions, or structural variants.

Training a retina-specific splice prediction tool did not lead to the identification of novel splice-altering variants in IRD patients. However, we anticipate that as more patients undergo WGS instead of exome sequencing to provide a genetic diagnosis, more pathogenic intronic and potentially splice-altering variants will be discovered. Moreover, it would be beneficial to train a splice prediction tool on pathogenic variants once enough training data is available rather than splice junctions in the genome, as this aligns with the primary application of most splice prediction tools.

## Code availability statement

All original code and models have been deposited at GitHub (https://github.com/cmbi/Retina-SpliceAI) and is publicly available.

## Acknowledgements

We thank the Radboud Genome Technology Center for infrastructural and computational support.

## References

1. Britten-Jones, A.C., Gocuk, S.A., Goh, K.L., Huq, A., Edwards, T.L., and Ayton, L.N. (2023). The Diagnostic Yield of Next Generation Sequencing in Inherited Retinal Diseases: A Systematic Review and Meta-analysis. Am J Ophthalmol 249, 57–73. 10.1016/J.AJO.2022.12.027.

2. Ben-Yosef, T. (2022). Inherited Retinal Diseases. Int J Mol Sci 23. 10.3390/IJMS232113467.

3. Wang, Z., and Burge, C.B. (2008). Splicing regulation: From a parts list of regulatory elements to an integrated splicing code. RNA 14, 802–813. 10.1261/RNA.876308.

4. Riolo, G., Cantara, S., and Ricci, C. (2021). What’s Wrong in a Jump? Prediction and Validation of Splice Site Variants. Methods Protoc 4. 10.3390/MPS4030062.

5. Jian, X., Boerwinkle, E., and Liu, X. (2014). In silico prediction of splice-altering single nucleotide variants in the human genome. Nucleic Acids Res 42, 13534. 10.1093/NAR/GKU1206.

6. Jaganathan, K., Kyriazopoulou Panagiotopoulou, S., McRae, J.F., Darbandi, S.F., Knowles, D., Li, Y.I., Kosmicki, J.A., Arbelaez, J., Cui, W., Schwartz, G.B., et al. (2019). Predicting Splicing from Primary Såequence with Deep Learning. Cell 176, 535-548.e24. 10.1016/j.cell.2018.12.015.

7. Cheng, J., Nguyen, T.Y.D., Cygan, K.J., Çelik, M.H., Fairbrother, W.G., Avsec, Ž., and Gagneur, J. (2019). MMSplice: Modular modeling improves the predictions of genetic variant effects on splicing. Genome Biol 20, 48. 10.1186/s13059-019-1653-z.

8. Desmet, F.O., Hamroun, D., Lalande, M., Collod-Bëroud, G., Claustres, M., and Béroud, C. (2009). Human Splicing Finder: An online bioinformatics tool to predict splicing signals. Nucleic Acids Res 37. 10.1093/nar/gkp215.

9. Yeo, G., and Burge, C.B. (2004). Maximum entropy modeling of short sequence motifs with applications to RNA splicing signals. In Journal of Computational Biology, pp. 377–394. 10.1089/1066527041410418.

10. Shapiro, M.B., and Senapathy, P. (1987). RNA splice junctions of different classes of eukaryotes: Sequence statistics and functional implications in gene expression. Nucleic Acids Res 15, 7155– 7174. 10.1093/nar/15.17.7155.

11. Rentzsch, P., Schubach, M., Shendure, J., and Kircher, M. (2021). CADD-Splice—improving genome-wide variant effect prediction using deep learning-derived splice scores. Genome Med 13, 31. 10.1186/s13073-021-00835-9.

12. Pertea, M. (2001). GeneSplicer: a new computational method for splice site prediction. Nucleic Acids Res 29, 1185–1190. 10.1093/nar/29.5.1185.

13. Reese, M.G. (1997). Improved splice site detection in Genie. In Journal of Computational Biology, pp. 311–323. 10.1089/cmb.1997.4.311.

14. Xiong, H.Y., Alipanahi, B., Lee, L.J., Bretschneider, H., Merico, D., Yuen, R.K.C., Hua, Y., Gueroussov, S., Najafabadi, H.S., Hughes, T.R., et al. (2015). The human splicing code reveals new insights into the genetic determinants of disease. Science (1979) 347, 1254806. 10.1126/science.1254806.

15. Naito, T. (2019). Predicting the impact of single nucleotide variants on splicing via sequence-based deep neural networks and genomic features. Hum Mutat 40, 1261–1269. 10.1002/humu.23794.

16. Zuallaert, J., Godin, F., Kim, M., Soete, A., Saeys, Y., and De Neve, W. (2018). Splicerover: Interpretable convolutional neural networks for improved splice site prediction. Bioinformatics 34, 4180–4188. 10.1093/bioinformatics/bty497.

17. Ellingford, J.M., Thomas, H.B., Rowlands, C., Arno, G., Beaman, G., Gomes-Silva, B., Campbell, C., Gossan, N., Hardcastle, C., Webb, K., et al. (2019). Functional and in-silico interrogation of rare genomic variants impacting RNA splicing for the diagnosis of genomic disorders. bioRxiv. 10.1101/781088.

18. Jian, X., Boerwinkle, E., and Liu, X. (2014). In silico tools for splicing defect prediction: A survey from the viewpoint of end users. Genetics in Medicine 16, 497–503. 10.1038/gim.2013.176.

19. Ohno, K., Takeda, J.I., and Masuda, A. (2018). Rules and tools to predict the splicing effects of exonic and intronic mutations. Wiley Interdiscip Rev RNA 9, 1451. 10.1002/wrna.1451.

20. Cheng, J., Çelik, M.H., Kundaje, A., and Gagneur, J. (2021). MTSplice predicts effects of genetic variants on tissue-specific splicing. Genome Biol 22, 94. 10.1186/s13059-021-02273-7.

21. Çelik, M.H., Wagner, N., Hölzlwimmer, F.R., Yépez, V.A., Mertes, C., Prokisch, H., and Gagneur, J. (2022). Aberrant splicing prediction across human tissues. bioRxiv, 2022.06.13.495326. 10.1101/2022.06.13.495326.

22. Zeng, T., and Li, Y.I. (2022). Predicting RNA splicing from DNA sequence using Pangolin. Genome Biol 23. 10.1186/s13059-022-02664-4.

23. Cao, H., Wu, J., Lam, S., Duan, R., Newnham, C., Molday, R.S., Graziotto, J.J., Pierce, E.A., and Hu, J. (2011). Temporal and Tissue Specific Regulation of RP-Associated Splicing Factor Genes PRPF3, PRPF31 and PRPC8—Implications in the Pathogenesis of RP. PLoS One 6. 10.1371/JOURNAL.PONE.0015860.

24. Liu, M.M., and Zack, D.J. (2013). Alternative splicing and retinal degeneration. Clin Genet 84, 142. 10.1111/CGE.12181.

25. Albert, S., Garanto, A., Sangermano, R., Khan, M., Bax, N.M., Hoyng, C.B., Zernant, J., Lee, W., Allikmets, R., Collin, R.W.J., et al. (2018). Identification and Rescue of Splice Defects Caused by Two Neighboring Deep-Intronic ABCA4 Mutations Underlying Stargardt Disease. Am J Hum Genet 102, 517–527. 10.1016/j.ajhg.2018.02.008.

26. Riepe, T. V., Khan, M., Roosing, S., Cremers, F.P.M., and ‘t Hoen, P.A.C. (2021). Benchmarking deep learning splice prediction tools using functional splice assays. Hum Mutat 42, 799–810. 10.1002/HUMU.24212.

27. Barbosa, P., Savisaar, R., Carmo-Fonseca, M., and Fonseca, A. (2023). Computational prediction of human deep intronic variation. bioRxiv, 2023.02.17.528928. 10.1101/2023.02.17.528928.

28. Smith, C., and Kitzman, J.O. (2023). Benchmarking splice variant prediction algorithms using massively parallel splicing assays. Genome Biology 2023 24:1 24, 1–22. 10.1186/S13059-023-03144-Z.

29. Pinelli, M., Carissimo, A., Cutillo, L., Lai, C.H., Mutarelli, M., Moretti, M.N., Singh, M.V., Karali, M., Carrella, D., Pizzo, M., et al. (2016). An atlas of gene expression and gene co-regulation in the human retina. Nucleic Acids Res 44, 5773–5784. 10.1093/NAR/GKW486.

30. Ratnapriya, R., Sosina, O.A., Starostik, M.R., Kwicklis, M., Kapphahn, R.J., Fritsche, L.G., Walton, A., Arvanitis, M., Gieser, L., Pietraszkiewicz, A., et al. (2019). Retinal transcriptome and eQTL analyses identify genes associated with age-related macular degeneration. Nat Genet 51, 606. 10.1038/S41588-019-0351-9.

31. Krueger, F., James, F., Ewels, P., Afyounian, E., and Schuster-Boeckler, B. (2023). FelixKrueger/TrimGalore: v0.6.10. Zenodo. 10.5281/zenodo.7598955.

32. Dobin, A., Davis, C.A., Schlesinger, F., Drenkow, J., Zaleski, C., Jha, S., Batut, P., Chaisson, M., and Gingeras, T.R. (2013). STAR: ultrafast universal RNA-seq aligner. Bioinformatics 29, 15–21. 10.1093/BIOINFORMATICS/BTS635.

33. Jun Kim, B., Choi, H., Jang, H., Lee, D., and Woo Kim, S. How to Use Dropout Correctly on Residual Networks with Batch Normalization.

34. Murphy, D., Cieply, B., Carstens, R., Ramamurthy, V., and Stoilov, P. (2016). The Musashi 1 Controls the Splicing of Photoreceptor-Specific Exons in the Vertebrate Retina. PLoS Genet 12, e1006256. 10.1371/journal.pgen.1006256.

35. Ciampi, L., Mantica, F., López-Blanch, L., Permanyer, J., Rodriguez-Marín, C., Zang, J., Cianferoni, D., Jiménez-Delgado, S., Bonnal, S., Miravet-Verde, S., et al. (2022). Specialization of the photoreceptor transcriptome by Srrm3-dependent microexons is required for outer segment maintenance and vision. Proc Natl Acad Sci U S A 119, e2117090119. 10.1073/pnas.2117090119.

36. Tapial, J., Ha, K.C.H., Sterne-Weiler, T., Gohr, A., Braunschweig, U., Hermoso-Pulido, A., Quesnel-Vallières, M., Permanyer, J., Sodaei, R., Marquez, Y., et al. (2017). An atlas of alternative splicing profiles and functional associations reveals new regulatory programs and genes that simultaneously express multiple major isoforms. Genome Res 27, 1759–1768. 10.1101/GR.220962.117.

37. Martín, G., Márquez, Y., Mantica, F., Duque, P., and Irimia, M. (2021). Alternative splicing landscapes in Arabidopsis thaliana across tissues and stress conditions highlight major functional differences with animals. Genome Biol 22. 10.1186/S13059-020-02258-Y.

38. Den Hollander, A.I., Koenekoop, R.K., Yzer, S., Lopez, I., Arends, M.L., Voesenek, K.E.J., Zonneveld, M.N., Strom, T.M., Meitinger, T., Brunner, H.G., et al. (2006). Mutations in the CEP290 (NPHP6) Gene Are a Frequent Cause of Leber Congenital Amaurosis. Am J Hum Genet 79, 556. 10.1086/507318.

39. Stone, E.M., Andorf, J.L., Whitmore, S.S., DeLuca, A.P., Giacalone, J.C., Streb, L.M., Braun, T.A., Mullins, R.F., Scheetz, T.E., Sheffield, V.C., et al. (2017). Clinically Focused Molecular Investigation of 1000 Consecutive Families with Inherited Retinal Disease. Ophthalmology 124, 1314. 10.1016/J.OPHTHA.2017.04.008.

40. Fadaie, Z., Whelan, L., Ben-Yosef, T., Dockery, A., Corradi, Z., Gilissen, C., Haer-Wigman, L., Corominas, J., Astuti, G.D.N., de Rooij, L., et al. (2021). Whole genome sequencing and in vitro splice assays reveal genetic causes for inherited retinal diseases. NPJ Genom Med 6. 10.1038/S41525-021-00261-1.

41. Reurink, J., Weisschuh, N., Garanto, A., Dockery, A., van den Born, L.I., Fajardy, I., Haer-Wigman, L., Kohl, S., Wissinger, B., Farrar, G.J., et al. (2023). Whole genome sequencing for USH2A-associated disease reveals several pathogenic deep-intronic variants that are amenable to splice correction. HGG Adv 4. 10.1016/J.XHGG.2023.100181.

42. de Bruijn, S.E., Rodenburg, K., Corominas, J., Ben-Yosef, T., Reurink, J., Kremer, H., Whelan, L., Plomp, A.S., Berger, W., Farrar, G.J., et al. (2023). Optical genome mapping and revisiting short-read genome sequencing data reveal previously overlooked structural variants disrupting retinal disease−associated genes. Genetics in Medicine 25, 100345. 10.1016/J.GIM.2022.11.013.

43. Li, H., and Durbin, R. (2009). Fast and accurate short read alignment with Burrows-Wheeler transform. Bioinformatics 25. 10.1093/bioinformatics/btp324.

44. Chen, X., Schulz-Trieglaff, O., Shaw, R., Barnes, B., Schlesinger, F., Källberg, M., Cox, A.J., Kruglyak, S., and Saunders, C.T. (2016). Manta: Rapid detection of structural variants and indels for germline and cancer sequencing applications. Bioinformatics 32. 10.1093/bioinformatics/btv710.

45. Roller, E., Ivakhno, S., Lee, S., Royce, T., and Tanner, S. (2016). Canvas: Versatile and scalable detection of copy number variants. Bioinformatics 32. 10.1093/bioinformatics/btw163.

46. Anna, A., and Monika, G. (2018). Splicing mutations in human genetic disorders: examples, detection, and confirmation. J Appl Genet 59, 253. 10.1007/S13353-018-0444-7.

47. Strauch, Y., Lord, J., Niranjan, M., and Baralle, D. (2022). CI-SpliceAI-Improving machine learning predictions of disease causing splicing variants using curated alternative splice sites. PLoS One 17, e0269159. 10.1371/JOURNAL.PONE.0269159.

48. de Sainte Agathe, J.M., Filser, M., Isidor, B., Besnard, T., Gueguen, P., Perrin, A., Van Goethem, C., Verebi, C., Masingue, M., Rendu, J., et al. (2023). SpliceAI-visual: a free online tool to improve SpliceAI splicing variant interpretation. Hum Genomics 17. 10.1186/S40246-023-00451-1.

49. Canson, D.M., Davidson, A.L., de la Hoya, M., Parsons, M.T., Glubb, D.M., Kondrashova, O., and Spurdle, A.B. (2023). SpliceAI-10k calculator for the prediction of pseudoexonization, intron retention, and exon deletion. Bioinformatics 39. 10.1093/BIOINFORMATICS/BTAD179.

50. Burnight, E.R., Fenner, B.J., Han, I.C., DeLuca, A.P., Whitmore, S.S., Bohrer, L.R., Andorf, J.L., Sohn, E.H., Mullins, R.F., Tucker, B.A., et al. (2023). Demonstration of the pathogenicity of a common non-exomic mutation in ABCA4 using iPSC-derived retinal organoids and retrospective clinical data. Hum Mol Genet. 10.1093/HMG/DDAD176.

51. Daiger, S., Rossiter, B., Greenberg, J., Christoffels, A., Hide, W., P Daiger, S., F. Rossiter, B., Greenberg, L.J., Christoffels, A., Hide, W., et al. (1998). Data services and software for identifying genes and mutations causing retinal degeneration. Invest Ophthalmol Vis Sci 39.

52. Waseem, N.H., Ortiz, A., Maubaret, C., Kennan, A., Humphries, P., Daiger, S., Webster, A., Jenkins, S., Bird, A.C., and Bhattacharya, S.S. (2006). Identification of Mutations in IMPDH1 in a Cohort of 96 Recessive RP Patients. Invest Ophthalmol Vis Sci 47, 5800–5800.

53. Bowne, S.J., Sullivan, L.S., Mortimer, S.E., Hedstrom, L., Zhu, J., Spellicy, C.J., Gire, A.I., Hughbanks-Wheaton, D., Birch, D.G., Lewis, R.A., et al. (2006). Spectrum and Frequency of Mutations in IMPDH1 Associated with Autosomal Dominant Retinitis Pigmentosa and Leber Congenital Amaurosis. Invest Ophthalmol Vis Sci 47, 34. 10.1167/IOVS.05-0868.

54. Dalla-Torre, H., Gonzalez, L., Mendoza-Revilla, J., Carranza, N.L., Grzywaczewski, A.H., Oteri, F., Dallago, C., Trop, E., Almeida, B.P. de, Sirelkhatim, H., et al. (2023). The Nucleotide Transformer: Building and Evaluating Robust Foundation Models for Human Genomics. bioRxiv, 2023.01.11.523679. 10.1101/2023.01.11.523679.

55. Leman, R., Parfait, B., Vidaud, D., Girodon, E., Pacot, L., Le Gac, G., Ka, C., Ferec, C., Fichou, Y., Quesnelle, C., et al. (2022). SPiP: Splicing Prediction Pipeline, a machine learning tool for massive detection of exonic and intronic variant effects on mRNA splicing. Hum Mutat 43, 2308. 10.1002/HUMU.24491.

56. Danis, D., Jacobsen, J.O.B., Carmody, L.C., Gargano, M.A., McMurry, J.A., Hegde, A., Haendel, M.A., Valentini, G., Smedley, D., and Robinson, P.N. (2021). Interpretable prioritization of splice variants in diagnostic next-generation sequencing. The American Journal of Human Genetics 108, 1564– 1577. 10.1016/J.AJHG.2021.06.014.

