## Supplementary Figure S1 for "Deep learning-based prediction of tissue-specific splice sites in the human neural retina"

### Supplementary Materials

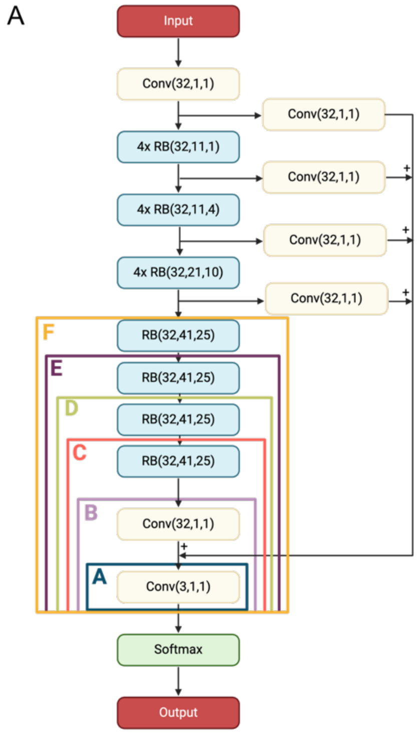


B

| Model | Retina PR-AUC | GTEx PR-AUC | Canonical PR-AUC | Number of trainable parameters |
|---|---|---|---|---|
| A | 0.74 | 0.87 | 0.96 | 163 |
| B | 0.74 | 0.86 | 0.97 | 1,219 |
| C | 0.75 | 0.86 | 0.97 | 85,379 |
| D | 0.75 | 0.86 | 0.97 | 169,539 |
| E | 0.76 | 0.86 | 0.96 | 253,699 |
| F | 0.76 | 0.86 | 0.96 | 337,859 |
| G | 0.80 | 0.84 | 0.96 | 696,803 |

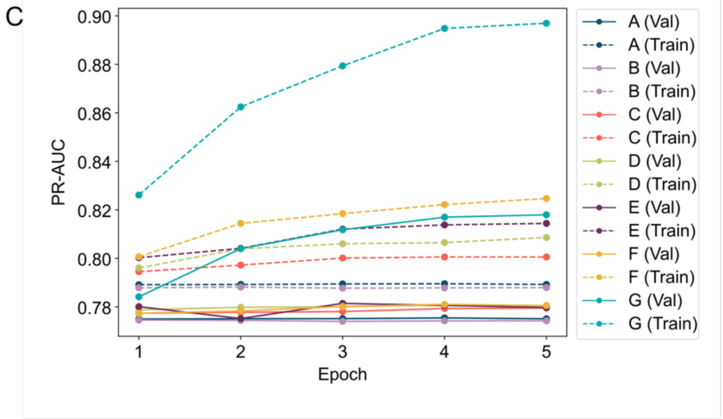


**Supplementary Figure S1: Overview of the transfer learning approach. (A)** Visualization of the weights that are fine-tuned for the different transfer learning approaches. For configuration G, all weights are fine-tuned. The SpliceAI model consists of residual blocks RB(N, W, D), where N, W, and D represent the number of convolutional kernels, window size, and dilation rate of each convolutional layer. Each RB block consists of a batch normalization layer, a ReLU activation layer, and a convolutional layer, repeated twice. **(B)** Number of trainable parameters and test PR-AUC values for each transfer learning model. **(C)** PR-AUC for the different transfer learning models at each training epoch.
